# Turning old foes into new allies – harnessing drainage canals for biodiversity conservation in desiccated novel ecosystems

**DOI:** 10.1101/2020.04.11.036897

**Authors:** Csaba Tölgyesi, Attila Torma, Zoltán Bátori, Jelena Šeat, Miloš Popović, Róbert Gallé, Nikolett Gallé-Szpisjak, László Erdős, Tamás Vinkó, András Kelemen, Péter Török

## Abstract

Drainage canals are ubiquitous components of agricultural landscapes worldwide. Although canals have greatly contributed to biodiversity loss by desiccating wetlands, they have recently attracted conservation attention due to their potential to function as refugia for native wetland-dependent species in intensively managed landscapes. However, their conservation role in complex landscapes comprising a mosaic of agricultural and desiccated semi-natural habitats, on which canals still pose a heavy burden, is unknown. Improved understanding of drainage canals and related biodiversity in these landscapes could help unlock their potential and support synergistic land management for nature conservation and water management.

We applied a multitaxon approach, including plants, butterflies, true bugs, spiders and birds, to (1) assess the conservation value of drainage canals in a heavily drained European lowland region, (2) to test landscape-level and local canal parameters for aiding prioritization among canal types, and (3) to propose a reconciliation-based management framework that suits the interest of all stakeholders.

We found that drainage canals concentrate more species across most taxa than adjacent semi-natural habitats, owing to the micro-environmental heterogeneity and the comparatively low management intensity in the canals. The species-concentrating capacity is particularly high in canals that traverse semi-natural habitats, although agricultural canals also support remarkable species diversity. However, agricultural canals are important dispersal corridors for invasive plants, which may negatively affect native species. Canal size has little effect on biodiversity but habitat stress is an important determinant. The higher the stress (due to sandiness and salinity), the higher is the added value of canals to landscape-wide biodiversity.

**Synthesis and applications:** We provide evidence that drainage canals can harbour surprisingly high levels of biodiversity and should therefore be recognized as important novel ecosystems with high conservation value, even within semi-natural habitats. Canals have previously been considered detrimental to nature conservation due to their association with loss of wetlands. However, by reducing water loss with reversible obstructions, controlling invasive species and applying specific conservation measures, they may be turned into conservation allies without compromising long-term interests of water management and agricultural land use.

## Introduction

Drainage and subsequent land cultivation has been a major threat to global wetland ecosystems for centuries (Herzon & Helenius, 2008; Blann et al., 2009; Davidson, 2016). In Europe, most lowland fens have been drained (Langheinrich et al., 2004; Hill et al., 2016); approximately 25% of the arable land of the United States is artificially drained (Herzon & Helenius, 2008), and immense wetlands have recently been drained in tropical Southeast Asia to gain land for agriculture (Aldhous, 2004). Canals, the main tools of drainage, are usually retained after the draining process and are regularly managed to sustain low and stable water balance in the cultivated landscape. Drainage canals are thus ubiquitous in heavily modified lowland agricultural landscapes worldwide (Shaw et al., 2015; Hill et al., 2016) and can form dense networks of interconnected artificial waterways. For instance, over 100,000 km of canals criss-cross the farmlands of the United Kingdom (Hill et al., 2016), and the total length of canals exceeds 300,000 km in the Netherlands (Blomqvist et al., 2003).

Despite being the main instrument of wetland loss, canals represent the only temporal continuity of wetland habitat in many drained landscapes (see Manhoudt et al., 2007 for the Netherlands), and are therefore refuges for organisms with high water demand (Chester & Robson, 2013; Harvolk et al., 2014). This paradoxical situation has led to the recognition that conservation value may be assigned to canals in agricultural landscapes, and therefore canals should be considered in conservation planning and agri-environmental schemes (Blomqvist et al., 2009; van Dijk et al., 2013).

Canals are often completely artificial with no functional or structural equivalent in the former natural wetlands; their vegetation and associated fauna can thus be considered as novel ecosystems (Hobbs et al., 2009). Management requirements for biodiversity in canals may be substantially different from those of natural wetlands, posing new challenges for conservation planners. Conventional management prescriptions of agri-environmental schemes have frequently been reported as ineffective (Blomqvist et al., 2009; van Dijk et al., 2013; Shaw et al., 2015). The main constraints for biodiversity in canals appear to be the high nutrient load, pollution with pesticides and herbicides, and the inappropriate intensity of bed management, including dredging and vegetation cutting (Herzon & Helenius, 2008; Blomqvist et al., 2009). However, when land managers have the tools and incentives to optimize management for biodiversity, canals can sustain populations of endangered species and high overall species richness, significantly increasing landscape-level conservation value (Manhoudt et al., 2007; Dorotovicova 2013; Tichanek & Tropek, 2015). Thus, canals have the potential to act as allies for biodiversity conservation in heavily transformed agricultural landscapes, despite that they were originally constructed to transform natural ecosystems.

The situation, however, is not so straightforward in moderately transformed landscapes where draining was not followed by intensive agriculture but wetlands turned into drier but still semi-natural habitats. In these landscapes, habitats surrounding the canals do not represent a hostile matrix but can also harbour significant biodiversity. The conservation role of these canals cannot be assessed in isolation, but only in conjunction with the surrounding habitats.

Studies on the biodiversity of canals that traverse habitats other than intensive arable fields are surprisingly scarce; papers dealing with such landscape configurations have mostly focussed on the hydrological, physical and chemical consequences of draining (e.g. Gasca-Tucker & Acreman, 2000; Gavin, 2003; Tiemeyer & Kahle, 2014). It is thus unknown whether these canals have an overall positive contribution to landscape level conservation value (i.e. local biodiversity maintenance in their bed vs. desiccating effects nearby), how to manage them in favour of biodiversity, or whether they should be maintained at all, if the opportunity to reverse-engineer them is an option.

Effective land management becomes more challenging in mosaic landscapes that are composed of both intensive agricultural fields and semi-natural habitats, interconnected with a network of drainage canals. This type of mosaic landscape may become more common in the future, due to increasing land abandonment and grassland restoration in formerly intensive agricultural landscapes of developed countries (Cramer et al., 2008; Valkó et al., 2016). Responsible land stewardship in these landscapes requires a complex understanding of the role of drainage canals harbouring novel ecosystems, and comprehensive guidelines must be developed for their management, in order to reconcile conservation purposes and immediate economic needs. At present, the scientific literature offers limited guidance for this endeavour but the emerging concepts of novel ecosystem management (Hobbs et al., 2009; Deák et al., 2020) and reconciliation ecology (Rosenzweig, 2003; Chapman et al., 2018) offer promising avenues. Considering these new fields of ecology, we aimed to understand the ecological role of the drainage canal network of a large, heavily drained European lowland region composed of a mosaic of intensive arable fields and semi-natural grasslands. Specifically, we aimed (1) to identify the extent to which drainage canals contribute to biodiversity conservation in regions rich in semi-natural habitats, (2) to test the effects of surrounding landscape matrix, channel size, soil substrate type, and reed and woody species abundance on the capacity of canals to sustain biodiversity, and (3) to propose reconciliation-based management guidelines which address the interests of different stakeholders (i.e. nature conservation and water management).

## Material and methods

### Study area

The study was carried out in the Danube-Tisza Interfluve, Central Hungary (Fig. 1). The climate is continental with a mean annual precipitation of 550-600 mm and temperature of 10-11°C (Tölgyesi et al., 2016). The soil substrate is diverse with mostly coarse sand in the central zone, saline loam along the bordering rivers and peaty loam (fen habitats) in between. Small isolated pockets of saline and fen areas also occur in the sandy central zone. This ca. 1 Mha lowland region used to be a mosaic of wetlands and drier habitats, but due to heavy draining in the middle of the 20th century, most wetlands vanished and were transformed into croplands, or gradually turned into grasslands (Biró et al., 2007; Ladányi et al., 2010). The promise of higher productivity land after draining proved to be mostly false, as natural ecosystems ceased to provide vital ecosystem services and productivity decreased in some high-lying arable fields due to severe groundwater decline. This landscape history is well reflected in the colloquial name of the main arterial drainage canal of the region: the “Cursed Channel” (Újházy & Biró, 2018).

**Figure 1.**
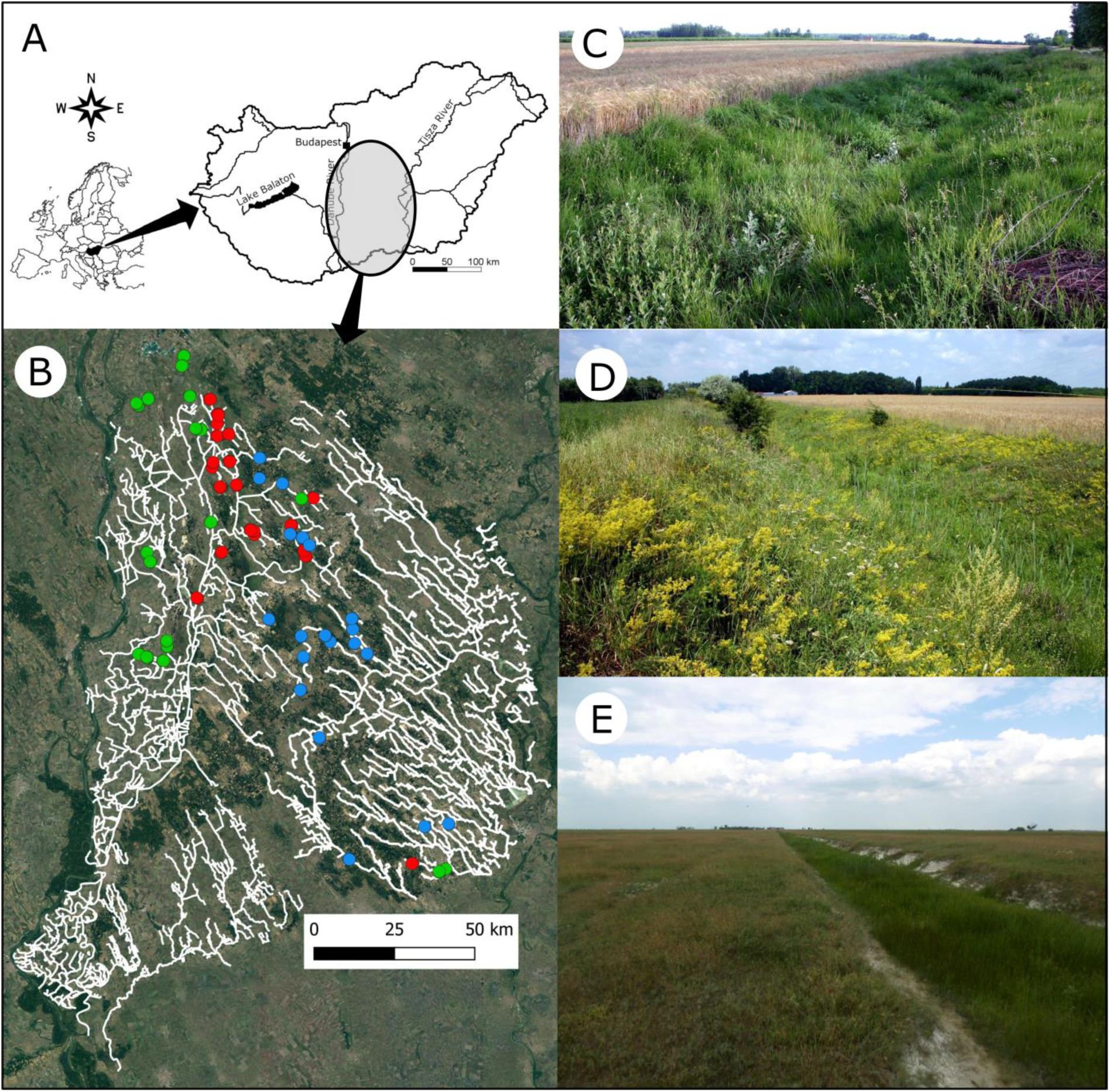
A: Location of the Danube-Tisza Interfluve in Hungary; B: registered drainage canal system of the region (white lines) and the position of the 60 studied canal sections; red: fen canals, green: saline canals, blue: sandy canals; C: small agricultural canal on fen substrate near Tázlár; D: large agricultural canal on sandy substrate near Kerekegyháza; E: small grassland canal on saline substrate near Dunapataj.

In the second half of the 20th century, regional aridification was further increased by climate change, specifically warming, prolonged droughts and an unfavourable redistribution of annual precipitation (Pongrácz et al., 2011), as well as increased groundwater extraction for irrigation and excessive afforestation (Biró et al., 2007; Tölgyesi et al. 2020). As a result, the water table greatly decreased (by up to 7 m in some localities; Ladányi et al., 2010) and the Food and Agriculture Organization of the United Nations classified some parts of the region as semi-desert (Tölgyesi et al., 2012). To date, counteractions have been limited to the filling up of some canals inside strict nature reserves and keeping sluices closed for longer periods than earlier, while the majority of canals are still functional and the other causes of aridification have not been addressed. The resulting environmental and biotic changes of water loss may have pushed the region over a tipping point into the realm of novel ecosystems (cf. Hobbs et al., 2006), in which the appropriate management of drainage canals may have a central role.

The total length of registered canals in the region is 4723 km (Fig. 1). Canals are infrequently managed by dredging (usually less than once a decade), reed cutting and shrub clearing. Mowing once a year and/or extensive grazing are the main management types of adjacent grasslands, but neither mowing nor grazing extends into the canals on a regular basis. Permanent water in the canals is nowadays rare; most contain water only in spring and after heavy rainy periods.

### Data collection

We selected sixty 200 m long drainage canal sections in the Danube-Tisza Interfluve, covering both agricultural and grassland canals (30 each), the three main substrate types, namely fens, saline areas and sandy areas (20 each) and both small and large canals (30 each), leading to five repetitions for each of the 12 category combinations (Fig. 1). Agricultural canals were fringed by at least 200 m wide croplands on both sides, while grassland canals were embedded in extensive grasslands that used to be wetter before draining. We identified two canal size classes: small ones with a depth of 0.7±0.2 m (mean±standard deviation) and a width of 3.6±1.4 m, and large ones with depth of 1.7±0.5 m and a width of 6.8±2.0 m. We also assessed the abundance of reed and woody vegetation by measuring their cumulative length along the canals and used them as additional variables to predict biodiversity. Canals dredged within the past ten years were not considered in the study.

To assess the biodiversity of canals, we applied a multi-taxon approach covering various functional groups, including primary producers (vascular plants), pollinating and herbivorous primary consumers (butterflies and true bugs, respectively), predators (spiders) and birds as representatives of large-bodied vertebrates. Vegetation was sampled in three ways during the summer of 2018. First, we compiled the total species pool of vascular plants in the 200-m sections (henceforth ‗gamma diversity‘) and second, recorded species in eight evenly spaced 1-m^2^ plots to capture plot-scale species density (henceforth ‗alpha diversity‘), making a total of 480 plots. Four plots were placed on the dry slopes of the bank and four plots into the bottom of the bed or adjacent to the bottom if water cover was too high. Third, we assessed the abundance of invasive plants. The measure was the cumulative length of invaded canal bank with a resolution of 1 m. Each bank was measured separately, leading to a maximum invasive plant abundance of 400 m.

Arthropod surveys were repeated three times in 2018: in spring (May), summer (July) and autumn (September) and were performed between 9 a.m. and 6 p.m. in dry, sunny weather without strong wind. We surveyed butterflies in transects along both canal banks. Each transect measured 200 m long and 2 m wide. The observer walked at a slow pace along the transect and counted all butterfly individuals seen 5 m in front of them (Nowicki et al., 2008). We sampled true bugs and spiders with sweep-netting. Each sample contained specimens from 25 sweeps prepared in a 25 m long transect, and we made four samples in every 200-m canal section. Samples per section were not pooled. We surveyed birds between 5 and 9 a.m. twice during the breeding season of 2019 (May and June). We scanned the area for ten minutes from an observation point adjacent to the canals, without disturbing the birds, then slowly walked by the canals to search for hiding individuals. We recorded every bird that landed on the vegetation or the surface of the canals; fly-bys were ignored.

We also selected three reference transects (two 2.5 m × 400 m transects for butterflies and a single 5 m × 200 m one for true bugs and spiders) parallel to every grassland canal, approximately 50 m from them. We performed all vegetation and arthropod surveys in the transects using the same protocol as in the canals. We did not attempt to prepare reference datasets for birds, as their density in the transects was low.

### Data analysis

For analysing the alpha and gamma diversity of the vegetation, we first discarded species of highly degraded and segetal communities (including invasive species) and retained only species of (semi-)natural grasslands and wetlands following the Flora Database of Hungary (Horváth et al., 1995). In line with the species pool hypothesis of (Zobel et al., 1998), we can expect different species pool sizes in different substrate types, making comparisons across substrates difficult. Therefore we standardized the species richness scores of canals to substrate specific average reference species richness scores, which yielded species excesses (or deficits, if negative), which we expressed as percentages. The use of species excesses was also beneficial because we were interested in the added conservation value of canals within a broader landscape. We used the following equation for the calculations:

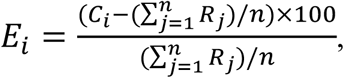

where *E*_*i*_ is the substrate specific species excess of the *i*th sampling unit of a canal (one of eight plots for alpha diversity or the total species count for gamma diversity), *C*_*i*_ is the species richness of this sampling unit, *R*_*j*_ is the species richness of the *j*th sampling unit of any of the reference transects belonging to the same substrate type as the canal, and *n* is the number of such reference sampling units.

Invasive plant species were either absent or very scarce in reference transects; therefore, we did not standardize their abundance in the canals but used the raw scores in subsequent analysis. For butterflies, true bugs and spiders, we applied the above method of standardization but the reference averages we used were specific to both substrate and season. In birds, we used the raw species richness scores for the analysis. Invasive or noxious pest species were not encountered among animal taxa; therefore, all species were retained for the analysis.

We applied a linear modelling approach to evaluate the biotic response variables (species excesses of plant and arthropod species richness, and raw invasive plant abundance and bird species richness). We had three categorical (landscape matrix, substrate and size) and two continuous (abundance of shrubs and reed) explanatory variables for plants and birds, while we included season as a fourth categorical explanatory variable in arthropods. We checked woody species and reed both as linear and quadratic terms and chose the one that resulted in a higher coefficient of determination (*R*^*2*^) in the models. If the quadratic term was chosen, we also identified the location of the hump or pit in the modelled response curve. We included canal identity as a random factor in models where multiple samples were collected in the canal sections. In birds, we used sampling occasion as a random variable. There was no indication of multicollinearity among the variables (generalized variance inflation factors ranged between 1.00 and 1.62); therefore all *a priori* variables were included in the final models.

In species excess type response variables, we also checked whether the mean score of each level of the categorical variables differed from the reference level (i.e. from the 0 score). For this analysis, we used reduced models including only one explanatory variable at a time and the random term of canal identity, if needed. In the case of arthropods, we also considered the repeated measures design, except when the seasons were tested for difference from the reference level.

Models were prepared in R environment (R Core Team, 2019) using the ‘lm’, ‘lmer’ (*lme4* package) or ‘glmer’ (*lme4* package, Poisson error term) functions depending on the data structures. We used the ‘MOStest’ function (*vegan* package) to identify the location of the hump or pit in models with quadratic terms, and the ‘fieller.MOStest’ function was applied to calculate 95% confidence intervals. Generalized variance inflation factors were calculated with the ‘vif’ function (*car* package). The significance of the explanatory variables was tested using the ‘Anova’ function (*car* package). Pairwise comparisons of the levels of substrate and season were performed with the ‘emmeans’ and ‘pairs’ functions (*emmeans* package).

## Results

### Plants

We recorded a total of 512 plant species in the study, but only 405 of these were of conservation interest (i.e. being wetland or grassland species); the remaining 107 species were weeds of highly degraded or segetal communities. Of the 405 species we included in the analysis, 380 occurred in the canals and 99 of these were not encountered in the reference grasslands. Species excess on the gamma level was significantly higher in grassland canals than in agricultural ones; large canals had higher species excesses than small ones and saline canals had higher values than fen canals. Compared to the habitat specific average reference levels, all levels of all categorical variables had significantly positive species excesses. Woody vegetation affected gamma diversity in a hump-shaped manner; the hump was located at 141.7 m but the upper limit of its confidence interval was beyond the upper limit of available woody abundance, rendering the prediction unreliable. The abundance of reed had no effect on the variation of the data (Fig. 2A, Table S1-3, Fig. S1A-B)

**Figure 2.**
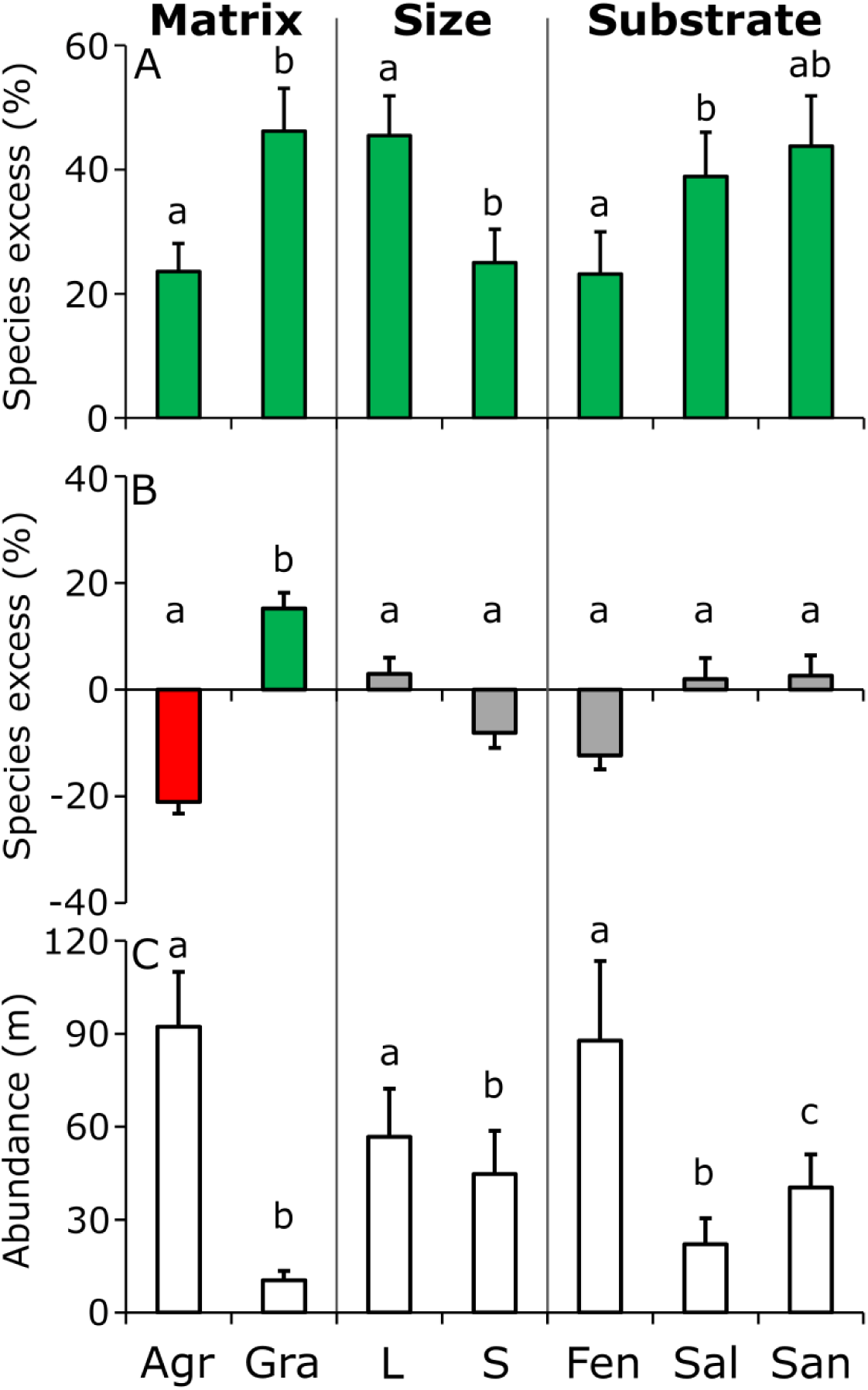
The effects of landscape matrix, canal size and substrate type on the species excesses of plants on the gamma (A) and the alpha (B) levels (i.e. considering total species counts of 200 m long canal sections and 1-m^2^ plots, respectively), and on the abundance of invasive species (C). Different lowercase letters within each canal parameter identify significantly different groups. Shading is used to denote differences from the reference level (i.e. the 0 score). Green and red indicate significantly positive and negative differences, respectively, while grey shading is used when no significant difference was detected from the reference level. Species excess is the proportional difference from the habitat specific reference averages expressed in per cents. No reference was generated for the abundance of invasives; therefore, no shading is used in subplot C. Agr: agricultural canals, Gra: grassland canals, L: large canals, S: small canals, Fen: fen substrate, Sal: saline substrate, San: sandy substrate. Whiskers show standard errors of the means.

We had more moderate results for species excesses on the alpha level, as it was not affected by canal size, substrate or the abundance of woody species and reed. However, agricultural canals had lower species excesses than grassland canals, and the former remained below the reference level (i.e. showed species deficit), while the latter slightly exceeded it. Mean values of small and large canals and canals of different substrate types did not differ from the reference levels (Fig. 2B, Table S1-3 and Fig. S1C-D).

We encountered several invasive plant species along the canal sections. Their cumulative abundance ranged between 0 m (absent) and 400 m (every metre of both banks invaded). The most abundant invasive species were *Asclepias syriaca*, *Solidago gigantea*, *S. canadensis* and *Aster lanceolatus* agg. Agricultural canals demonstrated a higher level of invasion than grassland canals, and large canals had more invasive species than small ones. Saline canals were the most intact, while fen and sandy canals had similar levels of invasion. The use of a quadratic term yielded better model fit for reed abundance and woody species abundance. The former had a significant effect on invasive species abundance with a pit at 69.4 m; however, the 95% confidence limits were beyond the available range, i.e. 0-200 m. The latter showed a hump at 70.3 m (95% confidence interval: 55.4 m – 97.8 m; Fig. 2C, Table S1-3 and Fig. S1E-F).

### Arthropods

We recorded 58 butterfly species (5962 adult individuals) in the study and 55 occurred in the canals, of which 19 were encountered only in the canals and not in the reference transects. Higher species excesses were found in grassland canals than in agricultural ones and the linear variant of woody abundance had a positive effect on butterfly species excess. Species excesses were higher in summer than in spring and autumn. Neither reed abundance, nor canal size, nor substrate had a significant effect on species excess. Compared to the substrate and season specific reference levels, only grassland canals and canals in summer had significantly positive species excesses (Fig. 3A, Table S1-3 and Fig. S2A-B).

**Figure 3.**
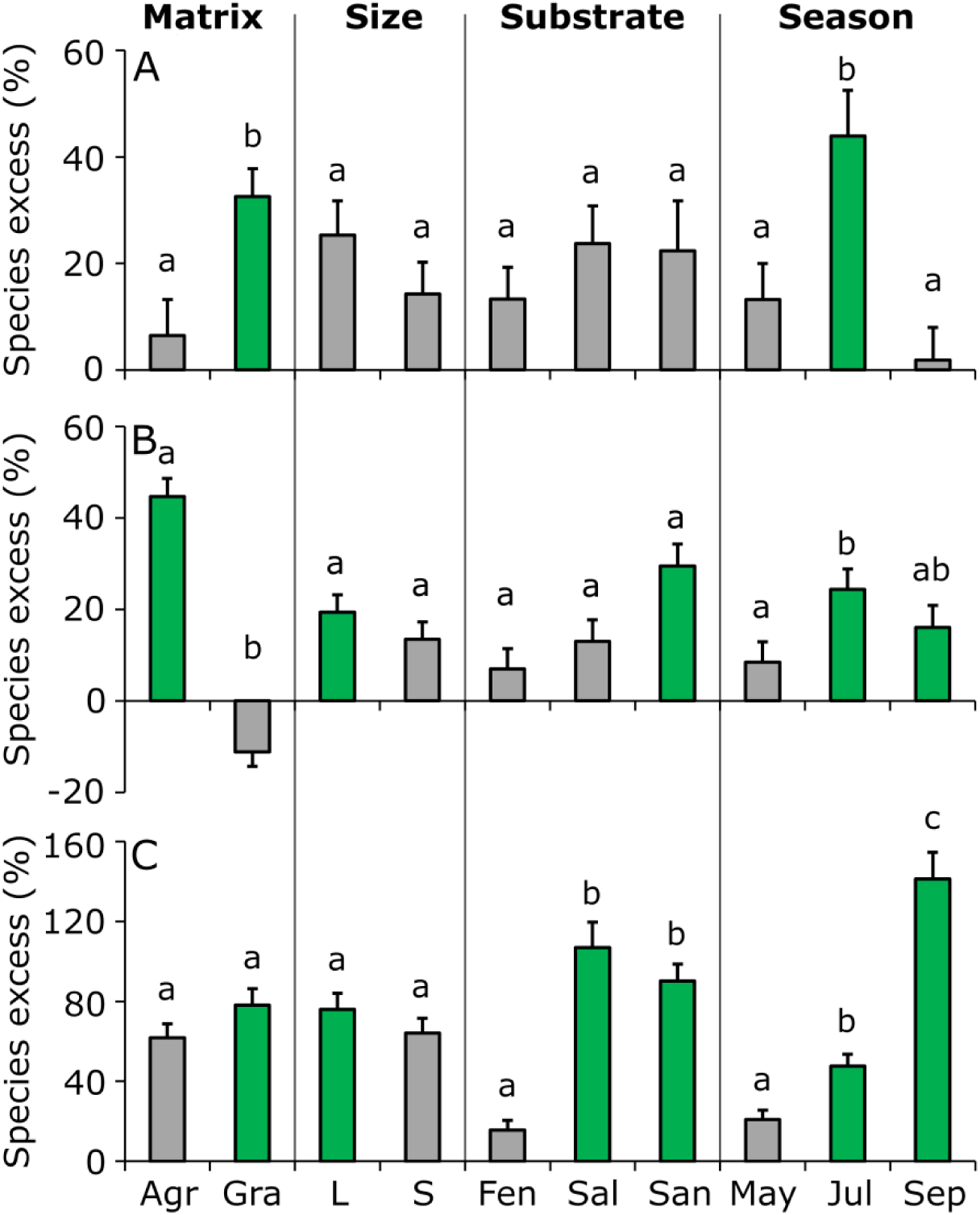
The effects of landscape matrix, canal size, substrate type and season on the species excesses of butterflies (A), true bugs (Heteroptera) (B) and spiders (C). Different lowercase letters within each canal parameter identify significantly different groups. Green shading indicates significantly positive difference from the reference level (i.e. the 0 score), while grey shading is used when no significant difference was detected from the reference level. Species excess is the proportional difference from the habitat and season specific reference averages. Agr: agricultural canals, Gra: grassland canals, L: large canals, S: small canals, Fen: fen substrate, Sal: saline substrate, San: sandy substrate. Whiskers show standard errors of the means.

We collected a total of 246 true bug species (30,012 adult individuals) in the study, and 219 occurred in the canals, of which 82 were collected exclusively there. Species excesses were affected only by matrix and season, with higher scores in agricultural canals than in grassland ones, and higher scores in summer than in spring. Compared to the reference levels, the statistics confirmed significant species excess in agricultural canals but not in grassland ones, in large canals but not in small ones, and on sandy substrate but not on fen or saline substrates. Species excess was highly positive in summer but significant difference was also confirmed for autumn data (Fig. 3B, Table S1-3 and Fig. S2C-D).

We recorded 134 spider species (6718 adult individuals) in the study; 114 occurred in the canals, of which 38 were found only there. Substrate and season had significant effects on species excesses, while matrix and size did not. Saline and sandy canals had higher species excesses than fen canals, and the excesses increased as the seasons progressed from spring through summer to autumn. Species excess was negatively affected by the linear variant of reed abundance, but woody cover had no detectable effect. Compared to the substrate and season specific reference levels, grassland canals had positive excesses but agricultural ones did not. Both large and small canals tended to have significantly positive scores, and among different substrate types, we could confirm positive species excesses for saline and sandy canals. Species excesses were significantly positive in summer and autumn but not in spring (Fig. 3C, Table S1-3 and Fig. S2E-F).

### Birds

We observed 38 bird species (892 individuals) during the study. We found that birds similarly frequented agricultural and natural canals and there was no difference among size classes and substrate types. However, the abundance of woody species and reed had significant effects and the models with quadratic terms fitted better than linear variants. Species excess had a hump along woody abundance at 113.6 m, although the upper confidence limit fell beyond the available range. In the case of reed, the hump itself was projected beyond the maximum of reed abundance (Fig. S2G-H, Fig. S3 and Table S1-3).

## Discussion

### Biodiversity of drainage canals

Drainage canals are ubiquitous components of managed lowland landscapes worldwide (Shaw et al., 2015; Hill et al., 2016). Although drainage is among the primary causes of the loss of the original biodiversity, there is growing evidence that canals can act as refuges for a variety of native species in desiccated and transformed landscapes (Chester & Robson, 2013; Golubovic et al., 2017; Torma et al., 2018), reinforcing the importance of moist microenvironments in the face of local and global environmental changes (Keppel et al., 2011; Mclaughlin et al., 2017). Our findings go one step further, as we show that canals not only function as secondary habitat for a certain subset of the native biodiversity, but they concentrate more species than adjacent semi-natural grasslands across a wide range of taxa. One reason for this diversity may be the micro-environmental heterogeneity offered by the canals (Stein et al., 2014). A wide moisture gradient is traversed from the dry top zone of the canal banks down to the bottom, enabling plant species with contrasting moisture demand to co-exist in close proximity, whereas the flat surrounding areas are characterized by more homogeneous environmental conditions. As a result, both grassland and agricultural canals could harbour more plant species of conservation interest (i.e. non-ruderal species) at the gamma level than semi-natural grasslands. Grassland canals had higher species richness even on the alpha level, despite the lack of annual management, which usually enhances the fine-scale co-existence of species (Klimek et al., 2007; Vadász et al., 2016).

The high diversity of plants, in turn, seems to cascade up to the level of primary consumers (butterflies and true bugs) and predatory arthropods (spiders). Nevertheless, some structural and functional features of the canals may have also contributed to the high arthropod richness we detected. Canals, which are local depressions in flat landscapes, can provide wind shelter for flying insects, including butterflies (Dover, 1996). Furthermore, canals provide overwintering opportunities for arthropods in the soil, litter and standing vegetation such as hollow stems, whereas soil disturbance in arable land impedes successful overwintering (Herzon & Helenius, 2008). Semi-natural grasslands and grassland canals may be similar in suitability for overwintering in the soil, but the management (i.e. mowing or grazing) of grasslands leaves little standing vegetation and litter into the winter for species that overwinter in these substrates. A variety of other non-cropped linear landscape elements, such as road verges, flower rich field margins or hedgerows have also been shown to be important for overwintering (Ramsden et al., 2015; Gallé et al. 2018), and canals are also likely to fulfil this function.

Arthropods greatly benefited from canals also in summer, as the highest species excesses were found in this period. Summers in the region are dry; thus canals can be important sources of water and fresh vegetation for true bugs and floral resources for butterflies. Furthermore, several studies emphasize that agricultural areas usually have “hunger periods” for arthropods in summer when there is a mismatch between resource demand and supply due to the synchronized phenology of cropped plants (Timberlake et al., 2019; Wintermantel et al., 2019). At the same time, canals are rich in resources throughout the vegetation period, including times when agricultural areas experience supply gaps. Although less commonly studied, this may also apply to managed grasslands. Grassland vegetation after being mown with powerful machinery provides little resource for either pollinators or other herbivory insects. Grassland canals are usually avoided during mowing, and thus represent continuity in food supply, similar to intentionally uncut vegetation strips in hay meadows (Buri et al., 2013; Kühne et al., 2015). Thus, canals are not just locally species-rich strips but are potentially important functional cornerstones of landscape-wide arthropod diversity.

Besides the local and landscape-level effects of canals on biodiversity, they may have consequences on even larger spatial scales. Canals often form continuous networks, overarching large regions and connecting isolated habitat fragments, similar to other linear landscape elements such as road verges and hedgerows (Vanniste et al., 2020). Canals, if longitudinally permeable for native species, can act as green corridors of dispersal, increasing regional connectivity and alleviating deficiencies of meta-population dynamics (van Geert et al., 2010; van Dijk et al., 2013), or can even act as conduits of climate change mediated range shifts, which would otherwise be hindered by extensive hostile areas, such as arable lands or exotic tree plantations (Saura et al., 2014; Robillard et al., 2015). Our findings, however, highlight that invasive species also use canals as dispersal corridors, especially in agricultural areas (see also Maheu-Giroux & de Blois 2007). As a result, canals can facilitate the invasion of otherwise intact and isolated habitats, and highly invaded canal sections may represent points of high resistance for native dispersal. Thus, controlling invasive species, particularly in agricultural canals, is a pressing issue that should be included in regional conservation strategies.

### Effects of canal parameters

We compared the biodiversity concentrating capacity of different canal types to aid prioritization among them, in order to channel conservation efforts where they are most needed. Although grassland canals proved to be more species rich for most taxa than agricultural ones, both types deserve our attention, as grassland ones exceed adjacent semi-natural grassland richness and agricultural canals represent the only conservation value in hostile agricultural landscapes. Large canals may be expected to have higher conservation value than small ones (Hill et al., 2016), but we found little evidence for this, as butterfly, true bug, spider and bird richness were not affected by canal size. Although plant species richness on the gamma level was higher in large canals, the abundance of invasive species was also higher in them, meaning that they are stronger conduits of plant invasion than small canals.

Conversely, substrate type provides more guidance for prioritization (see also Manhoudt et al., 2007). Fen canals proved to be the least valuable as (i) the rate of invasion was the highest in them, (ii) they had lower plant species excess than saline canals on the gamma level, and (iii) had lower spider species excess than the two other substrates. The reason for this may be twofold. Fen habitats provide the most benign conditions for plants as water availability is relatively high and stable, and no other stressors constrain plant life. These factors favour competitors, native and invasive ones alike, which can limit the number of co-existing species in canals (Blomqvist et al., 2003; Houlahan & Findlay, 2004) as compared to canals on other substrates. At the same time, adjacent fen grasslands are regularly managed to suppress competitors and sustain high species richness (Vadász et al., 2016), making habitat-specific reference species richness rather high.

The overall lower water supply and the more pronounced moisture gradient of canals on sand substrate (cf. Tölgyesi et al., 2016) may be the reason for the higher species excesses compared to fen canals. In saline canals, competitors may be further suppressed by the salt stress, and the gradients of salt and moisture can create diverse sets of micro-site conditions and associated species, similar to natural saline habitats with diverse micro-topography (Kelemen et al., 2013).

Regarding priority order, we conclude that landscape matrix and size are not decisive but substrate type should direct conservation efforts, as the higher the habitat stress (salinity in our model system), the higher the potential added conservation value. If this value cannot manifest on its own due to local conditions, these conditions should be identified and alleviated to ensure the full potential of canals in biodiversity concentration. In contrast, we cannot expect the same levels of added conservation value from canals under no-stress regimes; therefore, large scale conservation strategies should not target bringing the biodiversity level of these canals up to the same level as canals on other substrate types. This would be an unrealistic and/or an economically unfeasible aim.

Besides *a priori* canal parameters, we also tested the effects of parameters that can be modified by management (i.e. the abundance of woody species and reed). Moderate woody abundance had positive effects on biodiversity, while the effect tended to rebound at high abundances, leading to a hump for plants and birds and a pit for invasive species. Therefore some woody cover seems to have a positive effect on overall biodiversity. This finding was expectable, as sparse woody cover has been shown to introduce heterogeneity into micro-environmental conditions and vegetation structure (López-Pintor et al., 2006), both of which are known to boost biodiversity (Herzon & Helenius, 2008; Teleki et al., 2020).

The effect of reed abundance is more difficult to evaluate because it was contrasting among taxa (positive for birds but also positive for invasive plants, while negative for spiders). In fact, this was the only parameter that showed clear taxon specificity, suggesting that conservationists may need to choose which taxon to favour. Some authors have come to the conclusion that reed should regularly be cut along ditches for the benefit of biodiversity (e.g. Tichanek & Tropek, 2015) but these were mostly single-taxon studies, none of which considered birds. Since birds are declining rapidly in human-modified landscapes (Donald et al., 2001), the complete suppression of reed should be avoided to ensure that canals function as good quality refuge sites for them.

### Reconciliation between conservation and water management

Our findings highlight the high conservation value of drainage canals, but we need to emphasize that canals in their present form are not yet true allies in nature conservation. They still remove significant amounts of water from the surrounding landscape, contributing to water shortage both in semi-natural habitats (Ladányi et al., 2010; Pongrácz et al., 2011) and in agricultural production systems (Szinell et al., 1998). To make canals net positive contributors to conservation, the draining effect should be minimized while maintaining the canal profile with all the diverse microhabitats, flora and fauna. This can be achieved by introducing more sluices and semi-permanent obstructions, such as earthen plugs (see also Tichanek & Tropek, 2015). Canals that are permanently dry require no modification of the bed.

To date, conservationists have aimed to reverse-engineer entire canal sections in Hungary to fight against their draining effect (Valkó et al., 2017). This is extremely cost-intensive and, according to our results, the conservation benefit is questionable. Our recommendation is a more budget-friendly solution with no loss of biodiversity. If financial resources can be spared, they should be reallocated to identifying weak points of the canal network where the potential substrate-specific biodiversity concentrating effect is hindered by local factors, such as disturbance, pollution, and invasive infestation, and to designing customized local interventions to mitigate them.

We suppose that the introduction of more sluices and earthen obstructions would be more acceptable for water management authorities than removing the canals, since canals would thus remain available for reopening in case extreme water levels require. However, we discourage the application of regular dredging. In intensive agricultural areas of Western Europe, where canals experience high nutrient and pollutant loads, dredging is often recommended to suppress weedy species (Hill et al., 2016), but the canals in our study did not seem to require this intervention, which would probably promote invasive species. Furthermore, we encourage the retention of moderate amounts of woody plants and reed as they can increase the quality of canals as refuge sites for biodiversity. Implementation of these proposed guidelines would constitute a cost-effective, viable alternative to presently applied practices (Fig. 4), which can successfully reconcile nature conservation aims and water management in agricultural landscapes.

**Figure 4.**
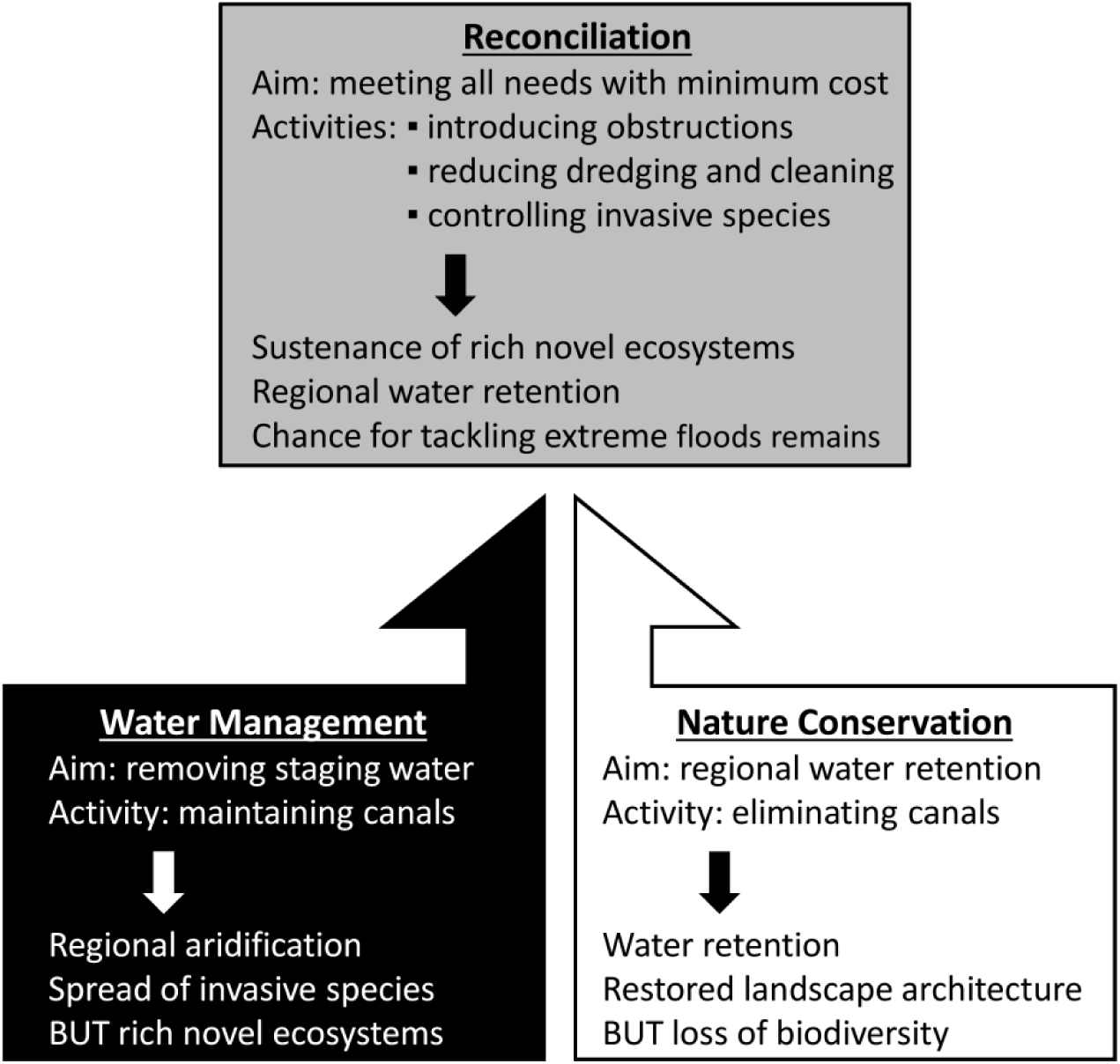
The main components of our management framework for drainage canals, showing the differing aims of stakeholders in the landscape and the related environmental impacts of their management (bottom left and right), and our proposed cost-effective management alternatives (top).

## Author Contributions

CT conceived the ideas and designed the methodologies, CT, JS, AT, RG, NG-S, MP, TV, ZB and AK participated in data collection, CT analysed the data, and all authors contributed to the writing of the manuscript.

## Acknowledgements

The support of the Hungarian Scientific Research Fund is greatly acknowledged (CT: 132131, ZB: K124796). The contribution of MP was funded by the Ministry of Education, Science and Technological Development of the Republic of Serbia (Project No. 173025). AK was supported by the Bolyai János Research Scholarship of the Hungarian Academy of Sciences, and JŠ received funding from the Stipendium Hungaricum Scholarship of the Tempus Public Foundation. The authors are grateful to Emmeline Topp for proofreading the manuscript.

## Data Availability

Data will be stored on the Dryad Digital Repository.

