## Supporting information for "Turning old foes into new allies – harnessing drainage canals for biodiversity conservation in desiccated novel ecosystems"

**Table S1** Coefficients of determination ($R^2$) of the four competing models of each taxon, in which the abundance of reed and the woody species were used as either linear or quadratic fixed factors. We chose the model with the highest value for further analysis (indicated with boldface). When models had identical values until the third decimal, we chose the linear version.

| | Plants - gamma | Plants - alpha | Invasive plants | Butterflies | True bugs | Spiders | Birds |
|---|---|---|---|---|---|---|---|
| Reed^1 + Woody^1 | 0.365 | 0.361 | 0.515 | **0.218** | 0.307 | **0.271** | 0.431 |
| Reed^2 + Woody^1 | 0.368 | 0.361 | 0.555 | 0.218 | 0.309 | 0.270 | 0.444 |
| Reed^1 + Woody^2 | 0.401 | **0.364** | 0.573 | 0.215 | 0.309 | 0.271 | 0.454 |
| Reed^2 + Woody^2 | **0.407** | 0.363 | **0.641** | 0.218 | **0.310** | 0.271 | **0.461** |

**Table S2** Test results of the fixed factors of the full models we prepared for the biotic variables. Pairwise comparisons of factor levels are shown for factors with more than two levels and with significant effect. The type of test statistics depends on the design of the models (for details see main text). Significant results (p<0.05) are indicated with boldface.

| | Plants - gamma | | Plants - alpha | | Invasive plants | | Butteflies | | True bugs | | Spiders | | Birds | |
|---|---|---|---|---|---|---|---|---|---|---|---|---|---|---|
| | *F* | *p* | *F* | *p* | *Chi*$^2$ | *p* | *F* | *p* | *F* | *p* | *F* | *p* | *Chi*$^2$ | *p* |
| Reed | 1.57 | 0.201 | 1.93 | 0.161 | **154.4** | **<0.001** | 3.64 | 0.058 | 3.61 | 0.164 | **5.35** | **0.021** | **22.92** | **<0.001** |
| Woody | **4.24** | **0.020** | 3.24 | 0.198 | **376.88** | **<0.001** | **8.92** | **0.003** | 2.02 | 0.363 | 0.53 | 0.468 | **26.94** | **<0.001** |
| Matrix | **11.82** | **0.001** | **27.14** | **<0.001** | **1575.3** | **<0.001** | **8.79** | **0.003** | **36.05** | **<0.001** | 0.36 | 0.551 | 0.79 | 0.375 |
| Size | **7.50** | **0.008** | 2.88 | 0.090 | **11.24** | **0.002** | 3.54 | 0.062 | 1.57 | 0.210 | 2.95 | 0.086 | 0.41 | 0.524 |
| Substrate | **4.99** | **0.010** | 6.31 | 0.053 | **575.3** | **<0.001** | 2.85 | 0.061 | 2.26 | 0.323 | **18.96** | **<0.001** | 0.51 | 0.774 |
| Season | | | . | . | | | **10.23** | **<0.001** | **9.87** | **0.007** | **136.21** | **<0.001** | | |
| | *t ratio* | *p* | *t ratio* | *p* | *t ratio* | | *t ratio* | *p* | *t ratio* | *p* | *t ratio* | *p* | *t ratio* | *p* |
| Fen-Saline | **-3.55** | **0.002** | . | . | **23.35** | **<0.001** | . | . | . | . | **-4.24** | **<0.001** | . | . |
| Fen-Sand | -1.03 | 0.562 | . | . | **18.28** | **<0.001** | . | . | . | . | **-2.57** | **0.034** | . | . |
| Saline-Sand | 2.30 | 0.065 | . | . | **-6.08** | **<0.001** | . | . | . | . | 1.52 | 0.289 | . | . |
| July-May | . | . | . | . | . | . | **-3.20** | **0.002** | **3.14** | **0.005** | **2.45** | **0.038** | . | . |
| July-Sept | . | . | . | . | . | . | **-4.37** | **<0.001** | 1.65 | 0.224 | **-8.66** | **<0.001** | . | . |
| May-Sept | . | . | . | . | . | . | -1.18 | 0.242 | -1.5 | 0.293 | **-11.11** | **<0.001** | . | . |

**Table S3** Test results of the reduced models in which we tested whether the levels of the categorical variables differ from the reference level. Biotic variables without reference levels were not analysed this way.

| | Plants - gamma | | Plants - alpha | | Butterflies | | True bugs | | Spiders | |
|---|---|---|---|---|---|---|---|---|---|---|
| | t | p | t | p | t | p | t | p | t | p |
| Matrix | | | | | | | | | | |
| • Agricultural | **4.07** | **<0.001** | **-4.09** | **<0.001** | 0.49 | 0.626 | **6.38** | **<0.001** | 1.65 | 0.102 |
| • Grassland | **8.07** | **<0.001** | **3.02** | **0.004** | **2.46** | **0.015** | -1.58 | 0.12 | **2.09** | **0.038** |
| Size | | | | | | | | | | |
| • Large | **7.74** | **<0.001** | 0.5 | 0.62 | 1.91 | 0.058 | **2.14** | **0.037** | **20.3** | **0.044** |
| • Small | **4.33** | **<0.001** | -1.34 | 0.184 | 1.08 | 0.282 | 1.5 | 0.139 | 1.72 | 0.088 |
| Substrate | | | | | | | | | | |
| • Fen | **3.23** | **0.002** | -1.68 | 0.098 | 0.96 | 0.338 | 0.49 | 0.625 | 0.41 | 0.680 |
| • Saline | **5.28** | **<0.001** | 0.27 | 0.789 | 1.7 | 0.09 | 0.9 | 0.369 | **2.80** | **0.006** |
| • Sand | **5.94** | **<0.001** | 0.39 | 0.698 | 1.61 | 0.109 | **2.86** | **0.006** | **2.37** | **0.019** |
| Season | | | | | | | | | | |
| • May | . | . | . | . | 1.84 | 0.068 | 1.22 | 0.22 | 1.8 | 0.072 |
| • July | . | . | . | . | **6.11** | **<0.001** | **3.58** | **<0.001** | **4.09** | **<0.001** |
| • September | . | . | . | . | 0.26 | 0.792 | **2.35** | **0.019** | **12.16** | **<0.001** |

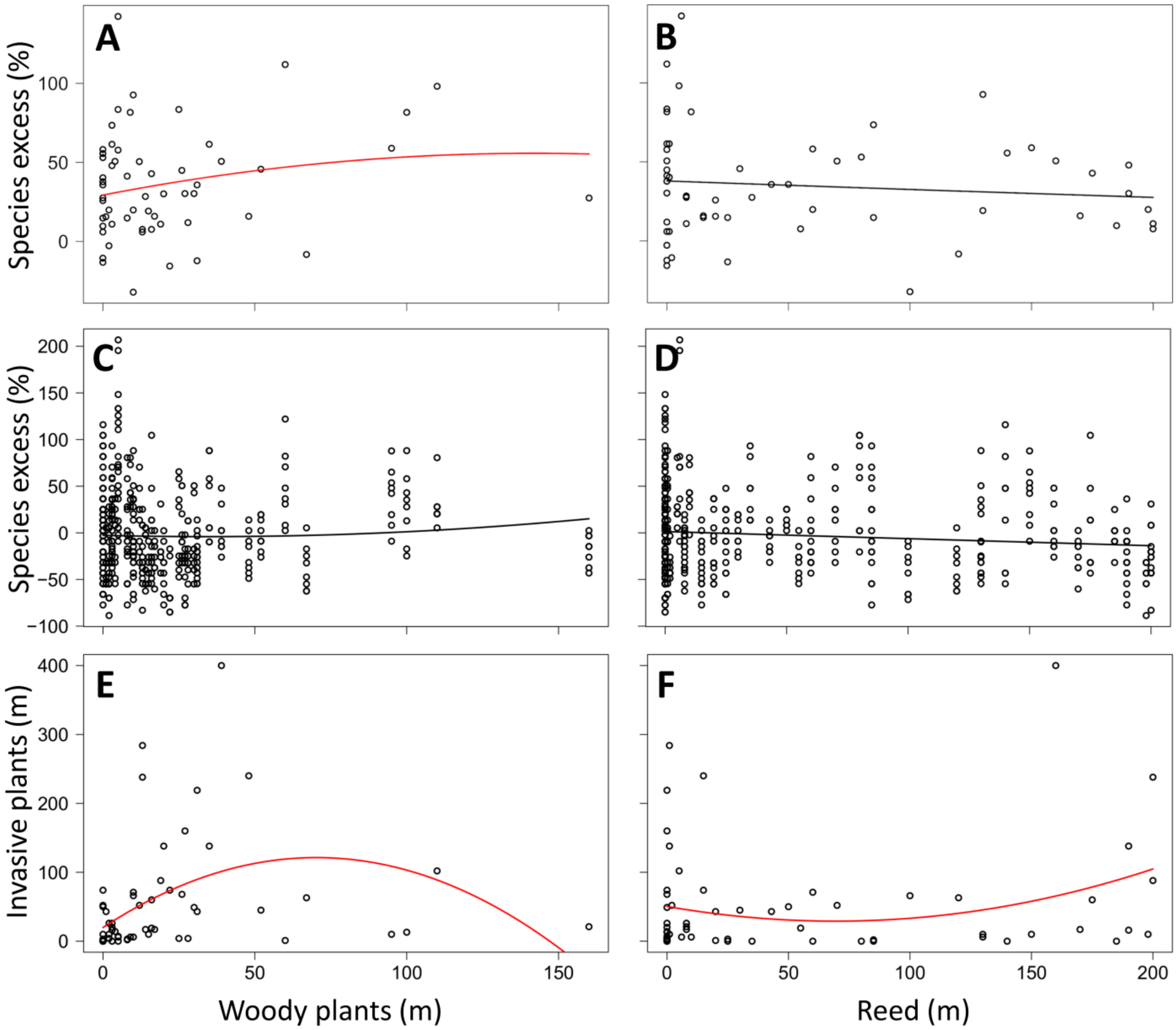


**Figure S1** Relationship between the botanical descriptors of drainage canals and the abundance of woody plants and reed. A and B: alpha diversity, C and D: gamma diversity, E and F: abundance of invasive species. Fitted models are either linear or quadratic; red ones explain significant proportion of the variation of data. The nested structure of the alpha diversity data (nested within site) is not reflected in the plots but considered in the models. ‘Species excess‘ is a proportional excess or deficit of species richness compared to habitat specific reference averages.

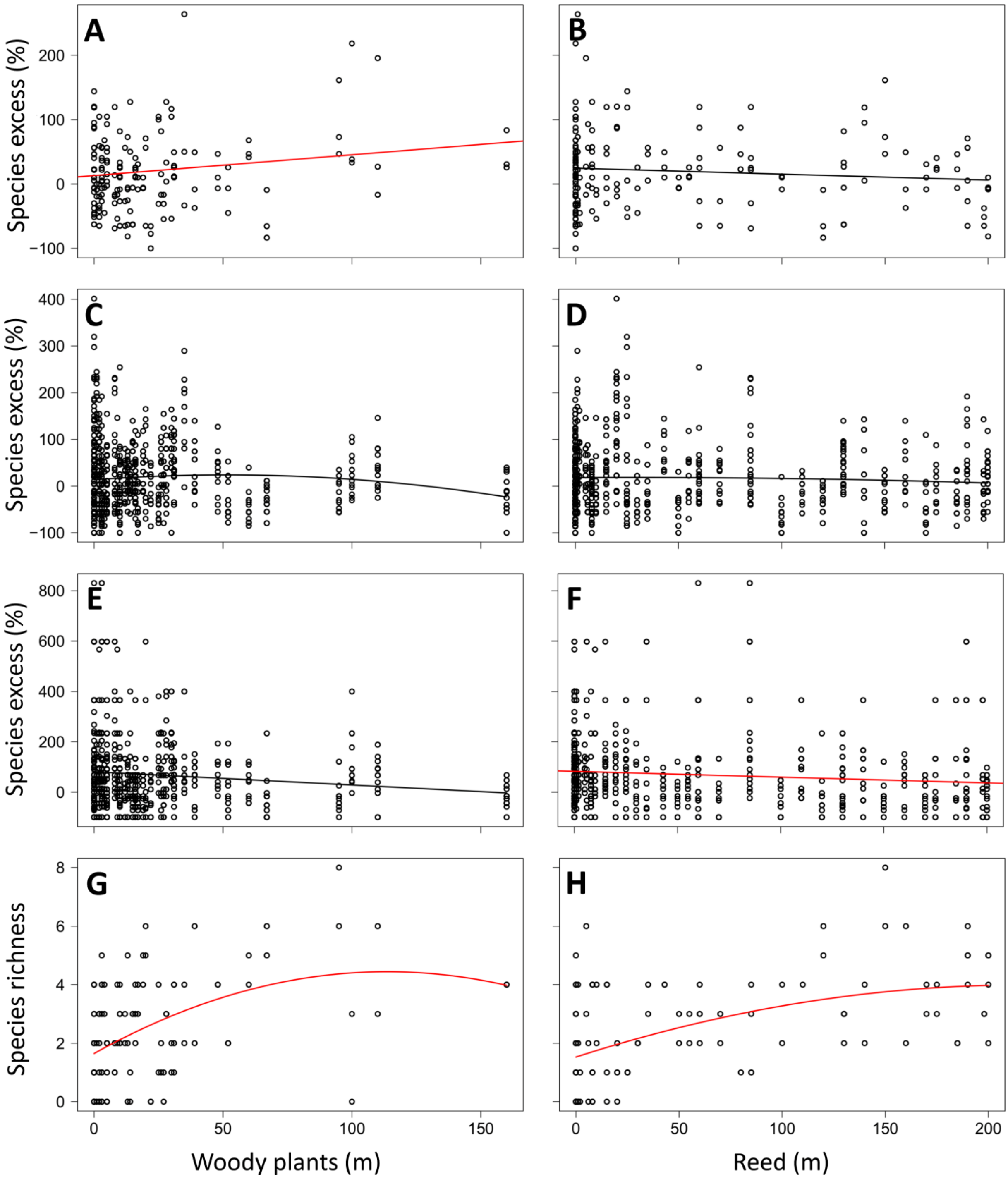


**Figure S2** Relationship between the zoological descriptors of drainage canals and the abundance of woody plants and reed. A and B: butterflies, C and D: true bugs, E and F: spiders, G and H: birds. Fitted models are either linear or quadratic; red ones explain significant proportion of the variation of data. The nested structure (nested within site) and repeated measures design of the data is not reflected in the plots but considered in the models. ‘Species excess‘ is a proportional excess or deficit of species richness compared to habitat specific reference averages.

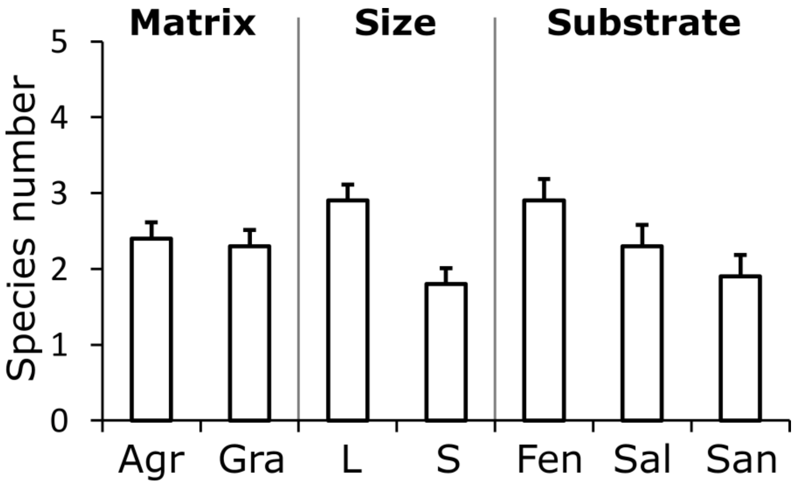


**Figure S3** Species richness of birds along the canal sections according to the studied categorical parameters. Agr: agricultural, Gra: grassland, L: large, S: small, Sal: saline, San: sandy canals. Whiskers show the standard error of the means. No significant differences were found among the levels of any parameter.
